# A highly discriminatory RNA strand-specific assay to facilitate analysis of the role of *cis*-acting elements in foot-and-mouth disease virus replication

**DOI:** 10.1101/2023.01.20.524889

**Authors:** Samuel J. Dobson, Joseph C. Ward, Morgan R. Herod, David J. Rowlands, Nicola J. Stonehouse

## Abstract

Foot-and-mouth-disease virus (FMDV), the etiological agent responsible for foot-and-mouth disease (FMD), is a member of the genus *Aphthovirus* within the *Picornavirus* family. In common with all picornaviruses, replication of the single-stranded positive-sense RNA genome involves synthesis of a negative-sense complementary strand that serves as a template for the synthesis of multiple positive-sense progeny strands. We have previously employed FMDV replicons to examine viral RNA and protein elements essential to replication, however, the factors affecting differential strand production remain unknown. Replicon-based systems require transfection of high levels of RNA, which can overload sensitive techniques such as qPCR preventing discrimination of specific strands. Here, we describe a method in which replicating RNA is labelled *in vivo* with 5-ethynyl uridine. The modified base is then linked to a biotin tag using click chemistry, facilitating purification of newly synthesised viral genomes or anti-genomes from input RNA. This selected RNA can then be amplified by strand-specific qPCR, thus enabling investigation of the consequences of defined mutations on the relative synthesis of negative-sense intermediate and positive-strand progeny RNAs. We apply this new approach to investigate the consequence of mutation of viral *cis*-acting replication elements and provide direct evidence for their roles in negative-strand synthesis.

## Introduction

The *Aphthovirus* foot-and-mouth disease virus (FMDV) is a member of the *Picornaviridae* family of single-stranded positive sense RNA viruses. FMDV is the etiological agent of foot-and-mouth disease (FMD) and comprises six serotypes; A, O, Asia 1, Southern African Territories (SAT) 1, SAT 2 and SAT 3, which are together responsible for endemic infection in large parts of Africa, Asia and the Middle East (1). Whilst FMD is rarely fatal, its consequences of reduced animal productivity, restriction of trade and slaughter of infected and at-risk animals result in severe economic losses (2, 3). The threat to maintenance of virus-free status in non-endemic regions is exacerbated by the high transmissibility of FMDV. Control measures following introduction of FMD into non-endemic regions, including strict movement restriction and mass culling, can result in costs of £billions (3–5). The antigenic diversity of FMDV adds to the challenges of disease control, with little cross-protection provided by strain-specific vaccines and no effective therapeutics currently available (2, 6–10). There is therefore a need to better understand the viral lifecycle in order to develop novel methods of control.

The genome of FMDV is approximately 8.4 kb and encodes a single polyprotein that is processed by viral proteases to generate the proteins required for genome replication and encapsidation. Primary cleavage of the polyprotein occurs at three positions to generate four products, the leader protease (L^pro^), the viral structural protein precursor P1-2A and the non-structural protein precursors P2 (2BC) and P3 (3AB_1,2,3_CD). Cleavage of L^pro^ occurs auto-catalytically, while release of the P1-2A precursor from the polyprotein occurs through a co-translational 2A-dependent ribosome skipping mechanism. The P1-2A region contains the viral structural proteins VP1, VP3 and VP0 (which is cleaved into VP2 and VP4 during virion maturation). Subsequent processing of the P2 and P3 precursors generates the viral non-structural proteins 2B, 2C and 3A, 3B_1,2+3_ (VPg), 3C protease (3C^pro^) and 3D polymerase (3D^pol^) (11–21). This processing is thought to be mediated by 3C^pro^ through alternative *cis-* and *trans-*pathways that appear to co-ordinate replication (17, 22, 23).

The single viral open reading frame (ORF) is flanked by untranslated regions (UTRs) at the 5’ and 3’ ends of the genome. The FMDV 5’ UTR is longest amongst known picornaviruses and contains several highly structured domains that are essential to the virus lifecycle (24–30). The S fragment is a ∼360 nucleotide hairpin-loop that occupies the same location (at the extreme 5’ end of the 5’ UTR) as the cloverleaf found in enteroviruses (31–34). It appears to be involved in immune modulation and viral replication via interactions with host and viral proteins (27, 35, 36). Adjacent to the S-fragment is a poly-C sequence of unknown function, followed by two to four RNA pseudoknots, which appear to be involved in assembly of infectious virions in addition to genome replication (24, 37–39). Adjacent to the pseudoknots is a small hairpin-loop known as the *cis*-acting replication element (*cre*) that in other picornaviruses is located within the ORF (25, 40). Whilst the location of the *cre* appears not important, it has an indispensable role in replication (25), acting as a template for the uridylylation of VPg by 3D^pol^ (41–43). VPg_p_U_p_U primes genome replication by 3D^pol^ (22, 44, 45). Finally, the internal ribosome entry site (IRES) is located immediately upstream of the ORF and is essential for initiation of translation (26, 29, 30, 46).

Picornavirus genome replication is initiated by the synthesis of a negative-sense copy of the infecting genome followed by assembly of a ‘replicative form’ (47–49). This complex comprises the negative strand template RNA and several nascent progeny positive strand RNAs together with 3D^pol^ and other viral and cellular proteins (50, 51). There is evidence that in the native state of the replicative form the nascent daughter strands are not collapsed onto the negative template strand but are only transiently associated at the site of transcription (50). The synthesis of both negative and positive RNA strands appears to be primed with VPg_p_U_p_U (41).

While it is well documented that the *cre* element is essential for viral replication, it is not firmly established whether this is required to produce VPgpUpU for the priming of both negative- and positive-sense RNA molecules during intra-cellular virus replication. Indeed, it has been reported that priming of negative-strand RNA synthesis can occur in a *cre*-independent manner during cell-free replication of poliovirus, possibly via the poly(A) tail (52–54). As mutation of the *cre* is lethal to progeny virus production, methods that facilitate the initiation of replicative events but are not reliant on infection (such as transfection of *in vitro* transcribed viral RNA) are useful for study of the initial steps in replication. Whilst assays to distinguish positive- and negative-strand RNAs have been developed for several viruses including FMDV, o’nyong-nyong virus, dengue virus, murine norovirus and chikungunya virus, their applications have so far been limited to studies involving infectious virus production (55–59).

Replicons are mini-genomes in which the structural genes are replaced with a reporter gene to facilitate the study of RNA replication independent of other aspects of the virus lifecycle (60). In addition to their application in the dissection of the molecular details of replication, replicons permit the study of viruses that require high-containment facilities, such as FMDV, at lower laboratory containment. However, because the delivery of RNA by transfection is an inefficient process, large quantities of *in vitro* transcribed RNA are used in order to ensure that sufficient cells are transfected. This can overload subsequent strand-specific assays. Here, we describe a strand-specific qPCR assay using FMDV replicons to determine the effects of mutations on the synthesis of both negative and positive strands. We have applied this method to determine the role of the *cre* element in the initiation of synthesis of both positive- and negative-strand RNAs.

## Materials and methods

### Cell lines and maintenance

Baby hamster kidney (BHK-21) cells were purchased from ATCC (LGC Standard) and were maintained in Dulbecco’s modified Eagle’s medium (DMEM) with glutamine, supplemented with 10% (v/v) foetal bovine serum (FBS), 50 U/mL penicillin and 50 μg/mL streptomycin. Growth medium and supplements were purchased from Sigma-Aldrich, Merck.

### *In vitro* transcription

The construction, linearisation and purification of FMDV replicon plasmids has been described previously (24, 60). Linear plasmids were transcribed *in vitro* using the T7 RiboMAX Express Large Scale RNA Production System (Promega) using ‘half sized’ reactions whereby 250 ng linear plasmid was added to a reaction mixture using half the volume suggested for all reagents in the manufacturer’s protocol. Reactions were incubated at 37°C for 1.5 hours prior to addition of 1 μL DNase I and further incubation at 37°C for 20 minutes. Resulting RNA was purified using the RNA Clean & Concentrator-25 kit (Zymo Research) prior to quantification by NanoDrop (Thermo Fisher) and confirmation of integrity by denaturing MOPS-formaldehyde gel electrophoresis.

### Replication assays

BHK-21 cells were seeded into 6-well plates at a density of 1×10^6^ cells/well and incubated for 16 hours. After incubation, cells were transfected with *in vitro* transcribed replicon RNA using Lipofectamine 2000 (Thermo Fisher). RNA (0.5 μg/cm^2^) and Lipofectamine 2000 (1 μg RNA:3 μL reagent) were added separately to 50 μL opti-MEM (Thermo Fisher) and incubated for 10 minutes at room temperature. RNA and reagent were mixed and incubated for 20 minutes at room temperature before diluting with phenol red free DMEM (Thermo Fisher) supplemented with 10% (v/v) FBS and GlutaMAX (Thermo Fisher) diluted to 1x. Cells were washed 1x using PBS, before addition of transfection mixture. Cells were imaged over time using the 10X objective of an Incucyte S3 live cell imaging system to detect phase and green fluorescence. Analysis was performed with the inbuilt Incucyte 2021A analysis suite using surface fit segmentation, threshold minimum (green calibrated units, GCU) 8.0, edge split sensitivity −30, minimum area filter 50 μm^2^ to determine green fluorescent cells. Following incubation for 6 hours, cells were washed 1x with PBS and extracted by addition of 1 mL TRI reagent (Merck) directly to the monolayer. Harvested cell extracts were stored at −20°C until further processing.

### RNA extraction and cDNA synthesis

Total RNA was TRI reagent extracted from BHK-21 cells using Phasemaker tubes (Invitrogen, Thermo Fisher) and associated manufacturer protocol. An additional 75% EtOH RNA pellet wash step was performed to ensure removal of trace contaminants prior to RNA solubilisation using nuclease-free H_2_O. Contaminating DNA was removed using the TURBO DNA-*free* kit (Thermo Fisher) following manufacturer guidelines. Total RNA purity and concentration was determined by NanoDrop.

cDNA synthesis was performed using SuperScript IV Reverse Transcriptase (Thermo Fisher) following manufacturer guidelines unless stated otherwise. Separate reactions were performed using 2 μM strand-specific reverse transcription (ssRT) primer (Table 1, PS-tag-RT or NS-tag-RT) or 50 μM oligo(dT)_20_. A total mass of 500 ng RNA was added to each reaction, together with dNTP mix containing 10 mM of each nucleotide (Promega). Combined reaction mixtures were incubated at 50°C for 10 minutes prior to inactivation at 85°C for 10 minutes.

**Table 1:**
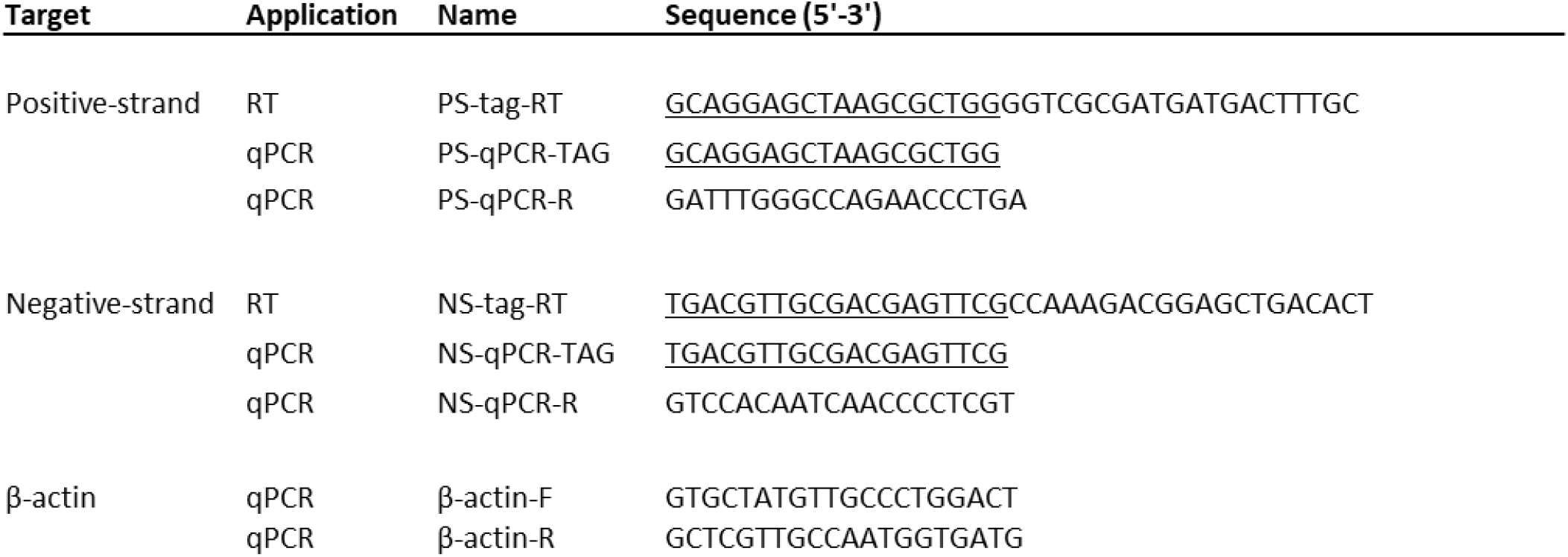
Primers used in RT and qPCR reactions. Underlined are the non-viral tag sequences for both the RT and qPCR primers.

### qPCR assays

cDNA synthesised from total RNA was diluted 100-fold in nuclease-free H_2_O unless otherwise stated. qPCR was performed using GoTaq qPCR Master Mix (Promega). Reaction mixes were made up in 10 μL total volume containing 500 nM forward and reverse primers (Table 1, PS-qPCR-tag & PS-qPCR-R or NS-qPCR-tag & NS-qPCR-R or β-actin-F & β-actin-R) and 2 μL diluted cDNA. qPCR conditions followed the manufacturer recommended fast cycling program with polymerase activation (95°C, 2 minutes) followed by 40 cycles (denaturation at 95°C, 3 seconds and annealing/extension at 60°C, 30 seconds) with melt curve analysis on a CFX Connect Real-Time PCR Detection System (Bio-Rad).

### Nascent RNA labelling assays

Nascent RNA labelling assays were performed using the Click-iT™ Nascent RNA Capture Kit (Thermo Fisher). Replication assays were performed as described previously in 6-well plates, with addition of 0.2 mM 5-ethynyl uridine (5-EU) diluted from a 200 mM stock, to transfection complex mixtures immediately prior to addition to cells. Cells were harvested 6 hours post transfection through the addition of 1 mL TRI reagent (Sigma-Aldrich, Merck) directly to the monolayer, with collected samples stored at −20°C until further processing. RNA was extracted as indicated above. Biotinylation of RNA by click reaction was performed following manufacturer guidelines using 10 μg total RNA and 0.5 mM biotin azide in a 50 μL reaction. RNA was precipitated in ammonium acetate overnight at −80°C, pelleted, washed and solubilised in nuclease-free H_2_O, as recommended. RNA was quantified by NanoDrop and 1 μg biotinylated RNA mixed with 12 μL Dynabeads® MyOne™ Streptavidin T1 magnetic beads. Following 30 minutes incubation at room temperature with agitation at 600 rpm, beads were washed for 3 minutes at room temperature with agitation at 600 rpm 5x using washer buffer 1 and 5x using wash buffer 2, with magnetic capture performed between washes. Bead mixtures were transferred to a new tube after every three washes to minimise tube contaminant carry over. Following the final wash, beads were resuspended in 36 μL wash buffer 2 and 12 μL slurry divided between three separate tubes for separate reverse transcription of positive-strand genomes, negative-strand genomes and total poly(A) RNA. Bead suspensions were heated at 70°C for 5 minutes prior to immediate addition of 17 μL master mix containing 15 μL NF-H_2_O, 1 μL 10 mM dNTPs and 1 μL 2 μM ssRT primer (Table 1) or 50 μM Oligo(dT)_20_. Mixes were cooled to room temperature with agitation at 600 rpm for 10 minutes before addition of 8 μL 5x SuperScript IV reaction buffer, 2 μL 100 mM DTT and 1 μL SuperScript IV enzyme to each reaction. Samples were incubated at 50°C for 1 hour with agitation at 600 rpm. Reaction mixtures were heated at 85°C for 10 minutes to inactivate reverse transcription and release cDNA from beads. Tubes were pulse centrifuged to pellet liquid prior to bead immobilisation using a magnetic rack and aspiration of cDNA. cDNA was diluted 100-fold for positive-strand reactions and 10-fold for negative-strand reactions. Oligo(dT)_20_ reactions were diluted accordingly. qPCR reactions were performed as previously described above with quantification using the ΔΔCq method. Samples in which no signal was detected were arbitrarily assigned a C_T_ value of 40 to permit quantification.

## Results

### Transfection with replicon RNA is not compatible with strand-specific discrimination by qPCR

Reverse genetics is a powerful tool for investigating the molecular biology of many viruses. However, a common drawback with many reverse genetics systems is the inefficient delivery of *in vitro* transcribed viral genomes, which can overwhelm assays deigned to dissect individual steps of viral lifecycles such as strand-specific genome synthesis. To overcome this restriction we have developed assays involving nascent RNA labelling, reverse transcription and strand-specific qPCR to examine the individual steps of negative- and/or positive-strand synthesis.

Our previously reported assays of replication utilised a FMDV replicon encoding the GFP reporter protein in place of the viral structural proteins (60, 61). Transfection of *in vitro* transcribed replicon RNA into cells allows measurement of a completed replication cycle via GFP expression prior to RNA extraction, cDNA synthesis and measurement of strand-specific specific RNA replication by qPCR (Figure 1A). Following transfection with *in vitro* transcribed pRep-ptGFP replicon RNA (WT or the replication-deficient 3D^GNN^) or yeast tRNA (as a control), cells were examined using an Incucyte S3 instrument to visualise phase contrast and green images (representative images Figure 1B) and the number of ptGFP positive cells determined using pre-defined analysis parameters (Figure 1C).

**Figure 1:**
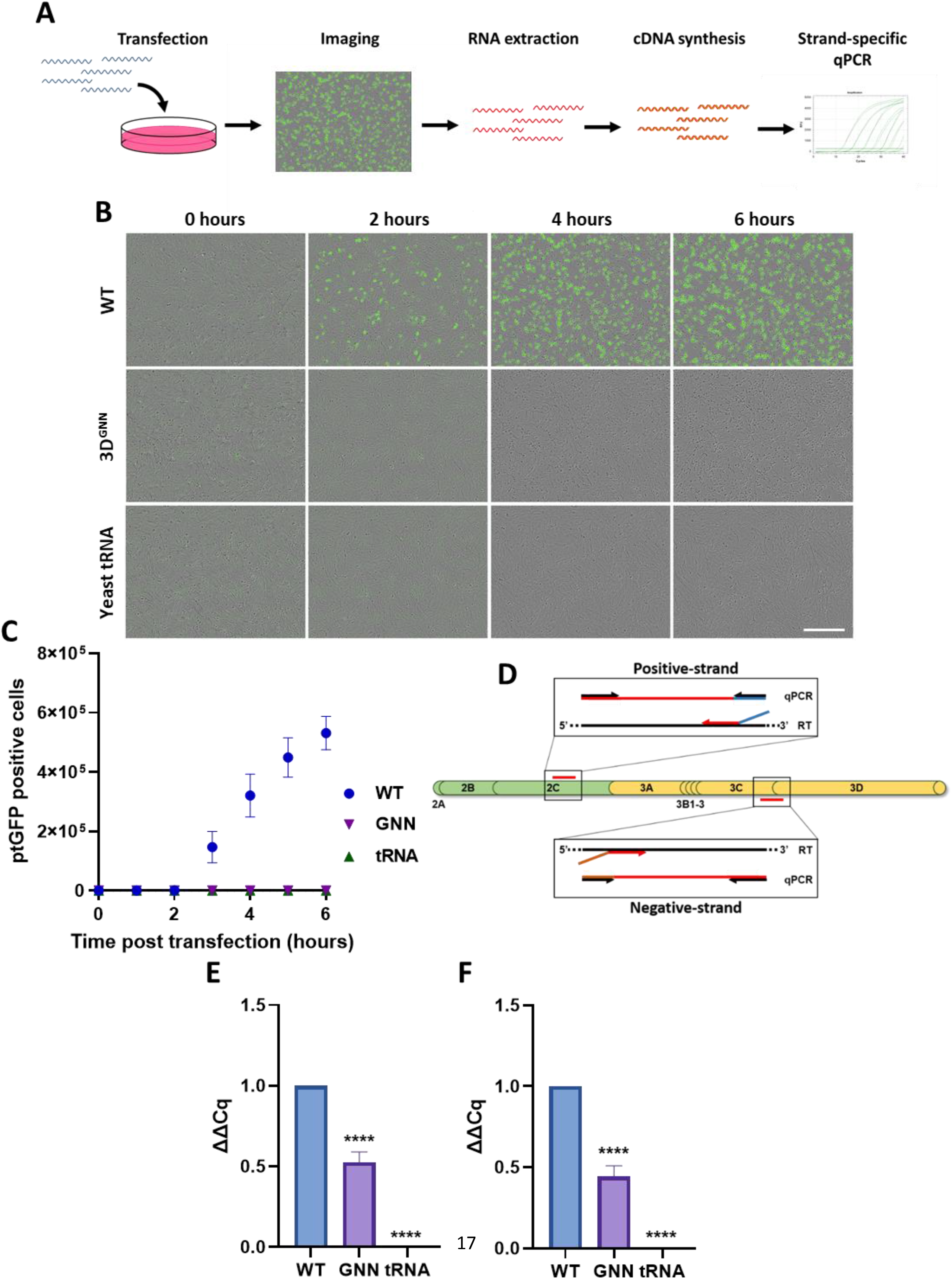
Transfection of FMDV replicon RNA is not compatible with distinguishing the synthesis of positive and negative strand molecules during replication. (A) Schematic overview of a replicon transfection assay with imaging prior to RNA extraction, cDNA synthesis and strand-specific qPCR. (B) Incucyte S3 imaging of BHK-21 cells transfected with WT and replication deficient 3D^GNN^ (GNN) FMDV pRep-ptGFP RNA and yeast tRNA (as a control), with phase contrast and green visualisation over time. Scale bar = 400 μm. (C) Detection over time of ptGFP following transfection with FMDV replicon RNA. (D) Schematic representation of primer binding regions with P2 and P3 genomic regions of FMDV for the cDNA synthesis and qPCR detection of positive- and negative-strands. RNA extracted from transfected cells was reverse transcribed using strand-specific primers prior to qPCR to detect positive-strand (E) and negative-strand (F) expression. Analysis was performed using the ΔΔCq method relative to WT. Statistical analysis was performed by one-way ANOVA with ** *P*-value = *<0*.*005* and **** *P-*value = *<0*.*0001*. n = 3.

Strand-specific primers with a non-viral tag sequence were designed for use during reverse transcription reactions to generate unique 5’ sequences providing specificity for each strand (Table 1) similar to published protocols (55–58). In order to minimise the potential for cross-contamination, primers were designed towards the 5’ and 3’ ends of the FMDV replicon genome for the positive-strand and negative-strand, respectively (Figure 1D). Reverse transcription using oligo(dT)_20_ was also performed for each sample to permit normalisation between samples by amplifying β-actin as a stably expressed reference gene. A primer complementary to the strand-specific non-viral tag and an internal primer complementary to the viral genome were designed for the strand-specific qPCR. Initial assays were performed comparing strand production between WT and 3D^GNN^ replicons. As anticipated, only a ≈2 fold-difference was observed between WT and 3D^GNN^ for both strands (Figure 1E&F), suggesting that the ability to demonstrate nascent viral RNA production was masked by the large quantity of input RNA necessary to efficiently initiate replication.

### Removal of input replicon genomes by nascent RNA labelling

We speculated that the inability of the strand-specific qPCR to distinguish between RNA isolated from cells transfected with either replication-competent or -incompetent replicon constructs was due to the large amount of input RNA necessary to initiate replication in the majority of the cells. To address this, we developed a method to specifically label and isolate newly synthesised RNA. To facilitate selection, transfected cells were incubated with an alkyne-modified uridine, 5-EU, from the time of transfection so that only cellular RNAs and viral genomes synthesised during replication incorporated the modified nucleotide (Figure 2A). Control assays indicated that addition of 0.2 mM 5-EU to the culture medium did not inhibit replication (Figure 2B).

**Figure 2:**
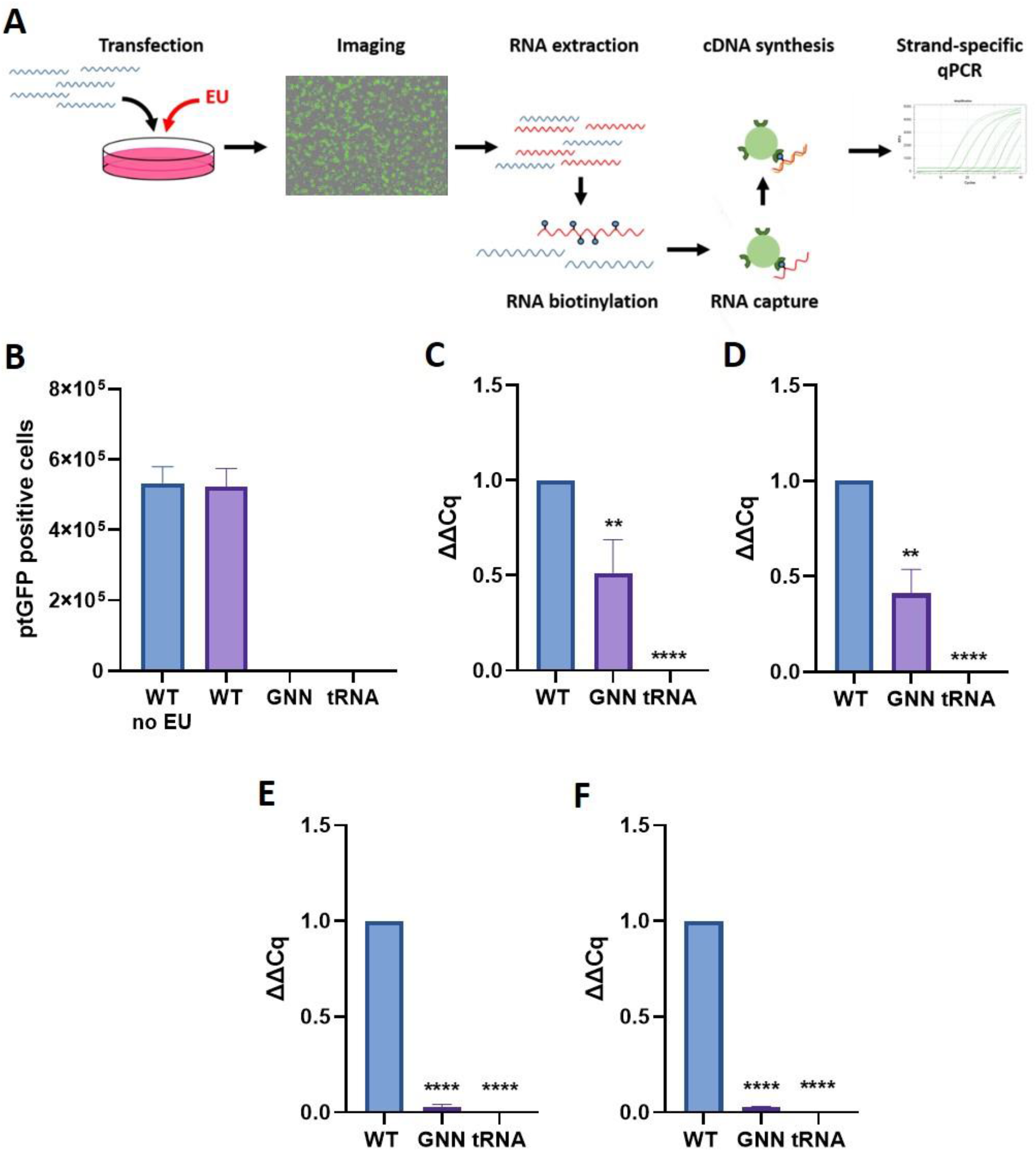
Removal of input replicon RNA by 5-EU labelling and RNA pulldown to select newly synthesised RNA. (A) Schematic representation of a 5-EU nascent RNA labelling assay. BHK cells were transfected with FMDV pRep-ptGFP RNA (WT and GNN), with addition of 0.2 mM 5-EU to label nascent RNA. An unlabelled control was also included (termed WT no EU). Cells were imaged using an Incucyte S3 instrument prior to harvesting of RNA 6 hours post transfection. 5-EU labelled RNA was biotinylated by click reaction and captured with streptavidin magnetic beads. Input RNA was removed by washing prior to on-bead cDNA synthesis and strand-specific qPCR. (B) ptGFP positive cells detected by Incucyte S3 imaging 6 hours post transfection to determine effect of 5-EU labelling on replication. Following RNA extraction, strand-specific qPCR to detect positive-strands (C) and negative-strands (D) prior to click reaction, was performed. Click reaction mediated biotinylation of nascent RNA, magnetic bead capture and on bead cDNA synthesis was performed prior to strand-specific assays to determine positive-strand (E) and negative-strand (F) expression. Analysis was performed using the ΔΔCq method relative to WT. Statistical analysis was performed by one-way ANOVA with ** *P*-value = *<0*.*005* and **** *P-*value = *<0*.*0001*. n = 3.

Following RNA extraction, strand-specific qPCR was performed to determine whether incorporation of 5-EU had any adverse effects upon the assay. These assays showed that detection of positive-strand and negative-strand genomes (Figure 2C&D, respectively) was comparable to that observed without labelling (Figure 1E&F). Copper-catalysed click reactions were performed to covalently link azide-modified biotin to the newly synthesised RNA. This acted as a bait for capture with streptavidin magnetic beads and removal by washing of input replicon RNA. Following capture and on-bead reverse transcription, analysis of the eluted cDNA confirmed that the method had efficiently removed replicon input RNA, with the 3D^GNN^ replicon signal reduced to 0.03 and 0.02 of WT for positive-strand and negative-strand, respectively (Figure 2E&F). This confirmed that labelling of newly synthesised RNA with 5-EU prior to biotin modification by click reaction and bead capture provided a method suitable for replicon-based investigation of differential viral strand synthesis.

### *Cre* is essential for negative-strand production

The *cre* is essential to replication of picornaviruses by acting as a template for uridylylation of VPg by 3D^pol^. There is also evidence that the poly(A) tail of poliovirus may act as a template for uridylylation in cell-free replication, albeit-it less efficiently (52–54). However, whether c*re* is essential for VPg uridylylation to prime the negative-strand intermediate during replication in cells is unknown. The role of the c*re* in strand synthesis was therefore investigated using replicons harbouring a complete c*re* deletion (Δ*cre*) or with a single nucleotide substitution of the functional AAACA motif (termed A1G, containing mutation of the first A nucleotide to G). Both of these mutations are known to prevent the complete virus lifecycle (25). Loss of replication competency was confirmed here for both mutants by replicon assay (Figure 3A). Strand-specific assays confirmed that the positive-strand could not be synthesised by either mutant replicon (Figure 3B). Negative-strand production was also ablated, showing that the c*re* is also essential during intracellular replication for production of the negative-strand (Figure 3C).

**Figure 3:**
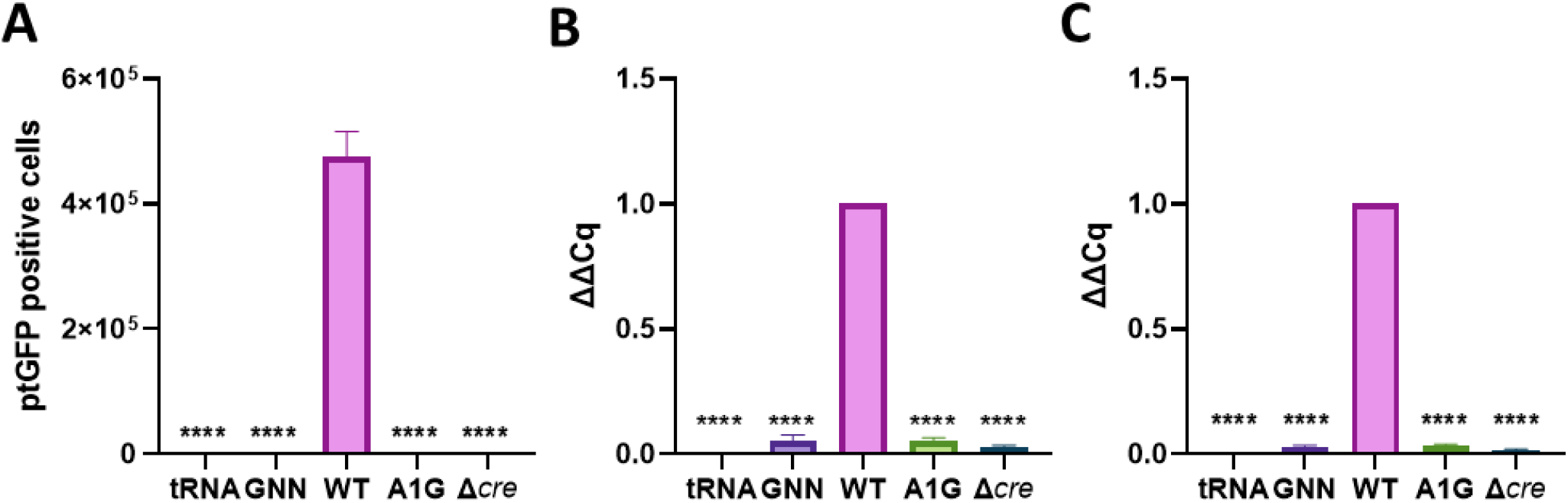
FMDV *cre* is essential to the production of negative-strand genomes. (A) BHK-21 cells were transfected with pRep-ptGFP constructs containing *cre* mutations A1G and Δ*cre* (alongside WT and GNN controls as described in Figure 1) with addition of 0.2 mM 5-EU to label nascent transcribed RNA. ptGFP positive cells were visualised 6 hours post-transfection using an Incucyte S3 instrument. Total RNA extracted 6 hours post transfection was biotinylated by click reaction, captured with streptavidin beads and reverse transcribed using strand-specific or oligo dT_(20)_ as a primer. cDNA was used in strand-specific qPCR assays to determine positive-strand (B) and negative-strand (C) expression. Analysis was performed using the ΔΔCq method relative to WT. Statistical analysis was performed by one-way ANOVA with **** *P-*value = *<0*.*0001*. n = 3.

## Discussion

The ability to reliably differentiate positive-strand and negative-strand RNAs is important for investigating details of the mechanism of picornavirus replication. This can be achieved using qPCR techniques, but the selection of strand-specific primers can be challenging. This problem is exacerbated when studying replicons, as reliable transfection requires a large quantity of input RNA. Despite these drawbacks, replicons facilitate investigations of replication that would otherwise be technically challenging, e.g. with non-recoverable or poorly replicating viruses, or to address issues regarding containment and safety. We therefore designed a strand-specific qPCR assay to distinguish synthesis of each strand separately and explored how mutation to the viral genome may restrict synthesis.

Initial assays identified the large quantity of input RNA required to ensure transfection of the majority of cells as a major restriction to demonstrating differences between replication-competent and -defective replicons (Figure 1). In the case of the FMDV replicon, the ≈ 7 kb genome relates to approximately 1.8 × 10^12^ genome copies/μg, and with up to 5 μg/well of replicon being transfected into cells, carry-over of input RNA was unsurprising. RNA self-priming, a consequence of high secondary structure at the 3’ terminus of RNA and reverse transcriptases lacking RNase H activity, has been reported previously as a source of contamination by complementary sequences in strand-specific assays (57, 58). In addition, residual fragments of plasmid DNA used as template for *in vitro* transcription of the transfecting RNA could theoretically also be a source of contaminating complementary sequences. However, contamination by any interfering sequences is eliminated by the specific selection of newly synthesised RNA as described here, thus providing an assay that can reliably distinguish complementary strands.

The essential role of the *cre* in picornavirus replication is well documented and was also confirmed here in replicon assays (Figure 3A) (36, 52–54, 62). It is possible that the poly(A) tail could template uridylylation of VPg (i.e. in addition to and independently of the c*re*, as reported in cell-free assays) (52–54). However, it was not established whether the c*re* is required for negative-strand synthesis in an intracellular assay. We therefore investigated the role of two c*re* mutations (A1G and Δ*cre*) during strand synthesis and found that neither positive-nor negative-strand RNAs were transcribed. Given that negative-strand intermediates were not produced, it is unsurprising that positive-strand production was also restricted. Due to a lack of negative-strand RNA it could not be established in this assay whether synthesis of new positive-strand genomes requires the *cre*. However, given evidence from the literature of the importance to positive-strand RNA synthesis, these findings would suggest that the c*re* is essential for efficient priming of replication by 3D^pol^ in the infected cell.

In conclusion, we present a method using 5-EU labelling and purification of nascent RNA prior to strand-specific qPCR that can be applied to determine the role of viral elements in transcription of positive- and negative-strands using replicons containing specific mutations. Here, this was applied to the known c*re* mutations A1G and Δc*re* and showed that neither are capable of producing a negative-strand intermediate. We will continue to probe other mutants in order to discover the role of viral elements that control viral replication of positive- and negative-strands.

## Acknowledgements

We are grateful to group members for their helpful comments on the project.

## Author contributions

The project was conceived by SJD, MRH, DJR and NJS. The experiments were carried out and the data analysed by SJD, JCW undertook mutagenesis. The manuscript was drafted by SJD and was edited and revised by all authors.

## References

1. Paton DJ, Di Nardo A, Knowles NJ, Wadsworth J, Pituco EM, Cosivi O, et al. The history of foot-and-mouth disease virus serotype C: the first known extinct serotype? Virus Evol. 2021;7(1):veab009.

2. Jamal SM, Belsham GJ. Foot-and-mouth disease: past, present and future. Vet Res. 2013;44:116.

3. Knight-Jones TJ, Rushton J. The economic impacts of foot and mouth disease - what are they, how big are they and where do they occur? Prev Vet Med. 2013;112(3-4):161–73.

4. Feng S, Patton M, Davis J. Market Impact of Foot-and-Mouth Disease Control Strategies: A UK Case Study. Front Vet Sci. 2017;4:129.

5. Thompson D, Muriel P, Russell D, Osborne P, Bromley A, Rowland M, et al. Economic costs of the foot and mouth disease outbreak in the United Kingdom in 2001. Rev Sci Tech. 2002;21(3):675–87.

6. Mahapatra M, Parida S. Foot and mouth disease vaccine strain selection: current approaches and future perspectives. Expert Rev Vaccines. 2018;17(7):577–91.

7. Park JH. Requirements for improved vaccines against foot-and-mouth disease epidemics. Clin Exp Vaccine Res. 2013;2(1):8–18.

8. Stenfeldt C, Eschbaumer M, Rekant SI, Pacheco JM, Smoliga GR, Hartwig EJ, et al. The Foot-and-Mouth Disease Carrier State Divergence in Cattle. J Virol. 2016;90(14):6344–64.

9. Dawe PS, Sorensen K, Ferris NP, Barnett IT, Armstrong RM, Knowles NJ. Experimental transmission of foot-and-mouth disease virus from carrier African buffalo (Syncerus caffer) to cattle in Zimbabwe. Vet Rec. 1994;134(9):211–5.

10. Dawe PS, Flanagan FO, Madekurozwa RL, Sorensen KJ, Anderson EC, Foggin CM, et al. Natural transmission of foot-and-mouth disease virus from African buffalo (Syncerus caffer) to cattle in a wildlife area of Zimbabwe. Vet Rec. 1994;134(10):230–2.

11. Gao Y, Sun SQ, Guo HC. Biological function of Foot-and-mouth disease virus non-structural proteins and non-coding elements. Virol J. 2016;13:107.

12. Roberts PJ, Belsham GJ. Identification of critical amino acids within the foot-and-mouth disease virus leader protein, a cysteine protease. Virology. 1995;213(1):140–6.

13. Vakharia VN, Devaney MA, Moore DM, Dunn JJ, Grubman MJ. Proteolytic processing of foot-and-mouth disease virus polyproteins expressed in a cell-free system from clone-derived transcripts. J Virol. 1987;61(10):3199–207.

14. Moffat K, Knox C, Howell G, Clark SJ, Yang H, Belsham GJ, et al. Inhibition of the secretory pathway by foot-and-mouth disease virus 2BC protein is reproduced by coexpression of 2B with 2C, and the site of inhibition is determined by the subcellular location of 2C. J Virol. 2007;81(3):1129–39.

15. Yost SA, Marcotrigiano J. Viral precursor polyproteins: keys of regulation from replication to maturation. Curr Opin Virol. 2013;3(2):137–42.

16. Ferrer-Orta C, Arias A, Agudo R, Perez-Luque R, Escarmis C, Domingo E, et al. The structure of a protein primer-polymerase complex in the initiation of genome replication. EMBO J. 2006;25(4):880–8.

17. Birtley JR, Knox SR, Jaulent AM, Brick P, Leatherbarrow RJ, Curry S. Crystal structure of foot- and-mouth disease virus 3C protease. New insights into catalytic mechanism and cleavage specificity. J Biol Chem. 2005;280(12):11520–7.

18. Grubman MJ, Zellner M, Bablanian G, Mason PW, Piccone ME. Identification of the active-site residues of the 3C proteinase of foot-and-mouth disease virus. Virology. 1995;213(2):581–9.

19. Forss S, Schaller H. A tandem repeat gene in a picornavirus. Nucleic Acids Res. 1982;10(20):6441–50.

20. King AM, Sangar DV, Harris TJ, Brown F. Heterogeneity of the genome-linked protein of foot- and-mouth disease virus. J Virol. 1980;34(3):627–34.

21. Gonzalez-Magaldi M, Martin-Acebes MA, Kremer L, Sobrino F. Membrane topology and cellular dynamics of foot-and-mouth disease virus 3A protein. PLoS One. 2014;9(9):e106685.

22. Ferrer-Orta C, Arias A, Perez-Luque R, Escarmis C, Domingo E, Verdaguer N. Structure of foot- and-mouth disease virus RNA-dependent RNA polymerase and its complex with a template-primer RNA. J Biol Chem. 2004;279(45):47212–21.

23. Herod MR, Gold S, Lasecka-Dykes L, Wright C, Ward JC, McLean TC, et al. Genetic economy in picornaviruses: Foot-and-mouth disease virus replication exploits alternative precursor cleavage pathways. PLoS Pathog. 2017;13(10):e1006666.

24. Ward JC, Lasecka-Dykes L, Neil C, Adeyemi OO, Gold S, McLean-Pell N, et al. The RNA pseudoknots in foot-and-mouth disease virus are dispensable for genome replication, but essential for the production of infectious virus. PLoS Pathog. 2022;18(6):e1010589.

25. Mason PW, Bezborodova SV, Henry TM. Identification and characterization of a cis-acting replication element (cre) adjacent to the internal ribosome entry site of foot-and-mouth disease virus. J Virol. 2002;76(19):9686–94.

26. Kanda T, Ozawa M, Tsukiyama-Kohara K. IRES-mediated translation of foot-and-mouth disease virus (FMDV) in cultured cells derived from FMDV-susceptible and -insusceptible animals. BMC Vet Res. 2016;12:66.

27. Kloc A, Diaz-San Segundo F, Schafer EA, Rai DK, Kenney M, de Los Santos T, et al. Foot-and-mouth disease virus 5’-terminal S fragment is required for replication and modulation of the innate immune response in host cells. Virology. 2017;512:132–43.

28. Carrillo C, Tulman ER, Delhon G, Lu Z, Carreno A, Vagnozzi A, et al. Comparative genomics of foot-and-mouth disease virus. J Virol. 2005;79(10):6487–504.

29. Lopez de Quinto S, Martinez-Salas E. Conserved structural motifs located in distal loops of aphthovirus internal ribosome entry site domain 3 are required for internal initiation of translation. J Virol. 1997;71(5):4171–5.

30. Lopez de Quinto S, Saiz M, de la Morena D, Sobrino F, Martinez-Salas E. IRES-driven translation is stimulated separately by the FMDV 3’-NCR and poly(A) sequences. Nucleic Acids Res. 2002;30(20):4398–405.

31. Xiang W, Harris KS, Alexander L, Wimmer E. Interaction between the 5’-terminal cloverleaf and 3AB/3CDpro of poliovirus is essential for RNA replication. J Virol. 1995;69(6):3658–67.

32. Gamarnik AV, Andino R. Interactions of viral protein 3CD and poly(rC) binding protein with the 5’ untranslated region of the poliovirus genome. J Virol. 2000;74(5):2219–26.

33. Harris KS, Xiang W, Alexander L, Lane WS, Paul AV, Wimmer E. Interaction of poliovirus polypeptide 3CDpro with the 5’ and 3’ termini of the poliovirus genome. Identification of viral and cellular cofactors needed for efficient binding. J Biol Chem. 1994;269(43):27004–14.

34. Yang Y, Rijnbrand R, Watowich S, Lemon SM. Genetic evidence for an interaction between a picornaviral cis-acting RNA replication element and 3CD protein. J Biol Chem. 2004;279(13):12659–67.

35. Yang F, Zhu Z, Cao W, Liu H, Wei T, Zheng M, et al. Genetic Determinants of Altered Virulence of Type O Foot-and-Mouth Disease Virus. J Virol. 2020;94(7).

36. Kloc A, Rai DK, Rieder E. The Roles of Picornavirus Untranslated Regions in Infection and Innate Immunity. Front Microbiol. 2018;9:485.

37. Zhu Z, Yang F, Cao W, Liu H, Zhang K, Tian H, et al. The Pseudoknot Region of the 5’ Untranslated Region Is a Determinant of Viral Tropism and Virulence of Foot-and-Mouth Disease Virus. J Virol. 2019;93(8).

38. Mohapatra JK, Pawar SS, Tosh C, Subramaniam S, Palsamy R, Sanyal A, et al. Genetic characterization of vaccine and field strains of serotype A foot-and-mouth disease virus from India. Acta Virol. 2011;55(4):349–52.

39. Escarmis C, Dopazo J, Davila M, Palma EL, Domingo E. Large deletions in the 5’-untranslated region of foot-and-mouth disease virus of serotype C. Virus Res. 1995;35(2):155–67.

40. Paul AV, Rieder E, Kim DW, van Boom JH, Wimmer E. Identification of an RNA hairpin in poliovirus RNA that serves as the primary template in the in vitro uridylylation of VPg. J Virol. 2000;74(22):10359–70.

41. Nayak A, Goodfellow IG, Belsham GJ. Factors required for the Uridylylation of the foot-and-mouth disease virus 3B1, 3B2, and 3B3 peptides by the RNA-dependent RNA polymerase (3Dpol) in vitro. J Virol. 2005;79(12):7698–706.

42. Schein CH, Ye M, Paul AV, Oberste MS, Chapman N, van der Heden van Noort GJ, et al. Sequence specificity for uridylylation of the viral peptide linked to the genome (VPg) of enteroviruses. Virology. 2015;484:80–5.

43. Paul AV, Yin J, Mugavero J, Rieder E, Liu Y, Wimmer E. A “slide-back” mechanism for the initiation of protein-primed RNA synthesis by the RNA polymerase of poliovirus. J Biol Chem. 2003;278(45):43951–60.

44. Paul AV, Peters J, Mugavero J, Yin J, van Boom JH, Wimmer E. Biochemical and genetic studies of the VPg uridylylation reaction catalyzed by the RNA polymerase of poliovirus. J Virol. 2003;77(2):891–904.

45. Paul AV, van Boom JH, Filippov D, Wimmer E. Protein-primed RNA synthesis by purified poliovirus RNA polymerase. Nature. 1998;393(6682):280–4.

46. Belsham GJ, Brangwyn JK. A region of the 5’ noncoding region of foot-and-mouth disease virus RNA directs efficient internal initiation of protein synthesis within cells: involvement with the role of L protease in translational control. J Virol. 1990;64(11):5389–95.

47. Bienz K, Egger D, Pfister T, Troxler M. Structural and functional characterization of the poliovirus replication complex. J Virol. 1992;66(5):2740–7.

48. Lundquist RE, Maizel JV, Jr. In vivo regulation of the poliovirus RNA polymerase. Virology. 1978;89(2):484–93.

49. Barton DJ, Flanegan JB. Synchronous replication of poliovirus RNA: initiation of negative-strand RNA synthesis requires the guanidine-inhibited activity of protein 2C. J Virol. 1997;71(11):8482–9.

50. Richards OC, Martin SC, Jense HG, Ehrenfeld E. Structure of poliovirus replicative intermediate RNA. Electron microscope analysis of RNA cross-linked in vivo with psoralen derivative. J Mol Biol. 1984;173(3):325–40.

51. Triantafilou K, Vakakis E, Kar S, Richer E, Evans GL, Triantafilou M. Visualisation of direct interaction of MDA5 and the dsRNA replicative intermediate form of positive strand RNA viruses. J Cell Sci. 2012;125(Pt 20):4761–9.

52. Goodfellow IG, Polacek C, Andino R, Evans DJ. The poliovirus 2C cis-acting replication element-mediated uridylylation of VPg is not required for synthesis of negative-sense genomes. J Gen Virol. 2003;84(Pt 9):2359–63.

53. Morasco BJ, Sharma N, Parilla J, Flanegan JB. Poliovirus cre(2C)-dependent synthesis of VPgpUpU is required for positive-but not negative-strand RNA synthesis. J Virol. 2003;77(9):5136–44.

54. Murray KE, Barton DJ. Poliovirus CRE-dependent VPg uridylylation is required for positive-strand RNA synthesis but not for negative-strand RNA synthesis. J Virol. 2003;77(8):4739–50.

55. Horsington J, Zhang Z. Analysis of foot-and-mouth disease virus replication using strand-specific quantitative RT-PCR. J Virol Methods. 2007;144(1-2):149–55.

56. Plaskon NE, Adelman ZN, Myles KM. Accurate strand-specific quantification of viral RNA. PLoS One. 2009;4(10):e7468.

57. Tuiskunen A, Leparc-Goffart I, Boubis L, Monteil V, Klingstrom J, Tolou HJ, et al. Self-priming of reverse transcriptase impairs strand-specific detection of dengue virus RNA. J Gen Virol. 2010;91(Pt 4):1019–27.

58. Vashist S, Urena L, Goodfellow I. Development of a strand specific real-time RT-qPCR assay for the detection and quantitation of murine norovirus RNA. J Virol Methods. 2012;184(1-2):69–76.

59. Muller M, Slivinski N, Todd E, Khalid H, Li R, Karwatka M, et al. Chikungunya virus requires cellular chloride channels for efficient genome replication. PLoS Negl Trop Dis. 2019;13(9):e0007703.

60. Tulloch F, Pathania U, Luke GA, Nicholson J, Stonehouse NJ, Rowlands DJ, et al. FMDV replicons encoding green fluorescent protein are replication competent. J Virol Methods. 2014;209:35–40.

61. Herod MR, Loundras EA, Ward JC, Tulloch F, Rowlands DJ, Stonehouse NJ. Employing transposon mutagenesis to investigate foot-and-mouth disease virus replication. J Gen Virol. 2015;96(12):3507–18.

62. Steil BP, Barton DJ. Cis-active RNA elements (CREs) and picornavirus RNA replication. Virus Res. 2009;139(2):240–52.

